# Identification of microbial communities and their removal efficiency of multiple pharmaceutical micropollutants combined in Membrane-Bioreactors

**DOI:** 10.1101/2023.04.11.536351

**Authors:** Marcel Suleiman, Francesca Demaria, Cristina Zimmardi, Boris Kolvenbach, Philippe Corvini

## Abstract

Pharmaceuticals are of concern to our planet and health as they can accumulate in the environment. The impact of these biologically active compounds on ecosystems is hard to predict and information on their biodegradation is necessary to establish sound risk assessment. Microbial communities are promising candidates for the biodegradation of pharmaceuticals such as ibuprofen, but little is known yet about their degradation-capacity of multiple micropollutants at higher concentrations (100 mg/L). In this work, microbial communities were cultivated in lab-scale Membrane Bioreactors (MBRs) exposed to increasing concentrations of a mixture of six micropollutants (ibuprofen, diclofenac, enalapril, caffeine, atenolol, paracetamol). Key players of biodegradation were identified using a combinatorial approach of 16S rRNA sequencing and analytics. Microbial community structure changed with increasing pharmaceutical intake (from 1 mg/L to 100 mg/L) and reached a steady-state during incubation for 7 weeks on 100 mg/L. HPLC analysis revealed a fluctuating but significant degradation (30-100%) of five pollutants (caffeine, paracetamol, ibuprofen, atenolol, enalapril) by an established and stable microbial community mainly composed of *Achromobacter*, *Cupriavidus*, *Pseudomonas* and *Leucobacter*. By using the microbial community from MBR1 as inoculum for further batch culture experiments on single micropollutants (400 mg/L substrate, respectively), different active microbial consortia were obtained for each single micropollutant. Microbial genera potentially responsible for degradation of the respective micropollutant were identified, i.e. *Pseudomonas* sp. and *Sphingobacterium* sp. for ibuprofen, caffeine and paracetamol, *Sphingomonas* sp. for atenolol, and *Klebsiella* sp. for enalapril. Our study demonstrates the feasibility of cultivating stable microbial communities capable of degrading simultaneously a mixture of highly concentrated pharmaceuticals in lab-scale MBRs and the identification of microbial genera potentially responsible for the degradation of specific pollutants.

**Graphical abstract:** 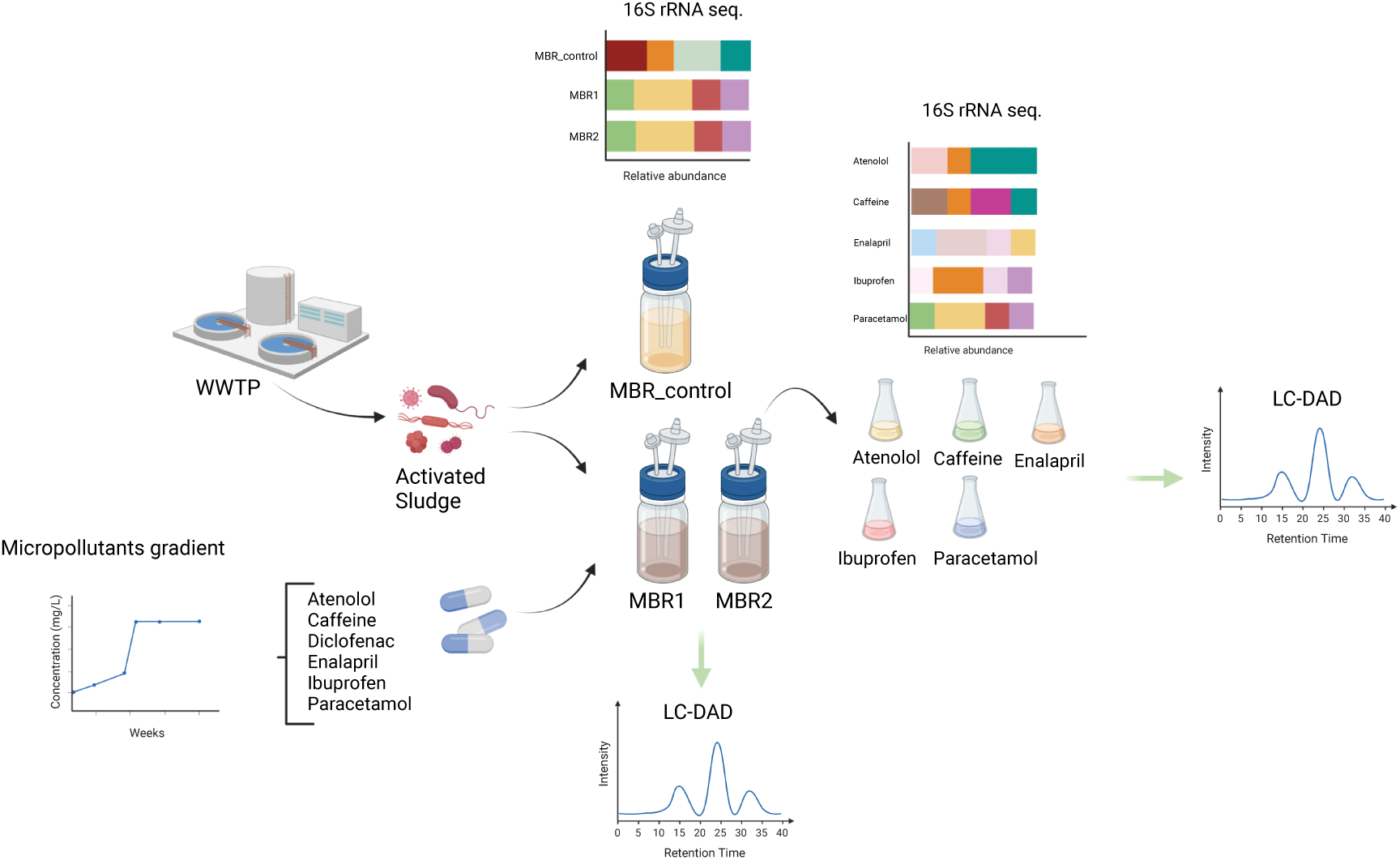

## Introduction

During the last decades, the production and consumption of pharmaceuticals increased significantly (Kristensen et al., 2016). Since many pharmaceuticals are not (totally) metabolized or assimilated in human and animal bodies, these biologically active compounds are partially eliminated in urine and feces before entering wastewater treatment plants (WWTP) in significant concentrations. Main sources of pharmaceutical micropollutants are hospitals, pharmaceutical industries, and animal farms (dos S. Grignet et al., 2022). A large portion of pharmaceutical residues in WWTP are, beside antibiotics, pain killers like ibuprofen (Buser et al., 1999), diclofenac (Vieno & Sillanpää, 2014), caffeine (Rigueto et al., 2020) and paracetamol (Wu et al., 2012), ß-blockers like atenolol (Salgado et al., 2013), and ACE inhibitors like enalapril (Chiarello et al., 2016). They are detected in ng/L to high µg/L range of concentration, depending on the location (Winker et al., 2008). Furthermore, the human consumption of pharmaceuticals is constantly increasing every year, resulting in high exposures and concentrations of these compounds in WWTP and the environment (dos S. Grignet et al., 2022).

Originally, WWTP were designed for the degradation of natural N-, P- and C-containing substrates, and increasing pharmaceutical intake poses a challenge for biodegradation of organic substances (BOD) (Khasawneh & Palaniandy, 2021). In some cases, the WWTP performance is not sufficient in terms of pollutants’ degradation and pharmaceutical contaminants can enter various environmental compartments (Dalahmeh et al., 2020). Consequently, pharmaceuticals are detected in rivers (Hughes et al., 2013), groundwater (Sui et al., 2015) and soils (Thiele-Bruhn, 2003). Little is known about the long-term impact of these biologically active contaminants on ecosystems and human health. Performing wastewater treatment plants are therefore crucial for the elimination of these micropollutants. Membrane bioreactors (MBRs) have a large potential for wastewater treatment, combining biodegradation with membrane filtration systems enabling biomass retention (Al-Asheh et al., 2021). In MBRs operated with infinite retention time no excess sludge is taken and evolutionary processes may even improve the microbial functionality concerning biodegradation (Zheng et al., 2019; Zhuang et al., 2016). Microbial communities are key players of the MBR concept, and their structure and performance are crucial for the efficient removal of pharmaceutical pollutants and the release of non-toxic effluents. Therefore, there is a strong need to identify promising bacterial communities for further development of microbial formulations to be used for bioaugmentation purpose. In this study, MBRs were operated with increasing concentration (1-100 mg/L) of a mixture consisting of six pharmaceutical pollutants (atenolol, caffeine, diclofenac, enalapril, ibuprofen, paracetamol). The choice of applying a mixture of pharmaceuticals was motivated by previous studies showing that multiple drivers can affect microbial communities differently (Orr et al., 2022; Suleiman et al., 2022). In order to analyze key players for the degradation of each pharmaceutical, MBR communities were transferred to batch cultures incubated with one single micropollutant. The potential of microbial communities to remove highly concentrated pharmaceutical was analyzed and microbial genera involved in degradation were identified for each pharmaceutical.

## Material and methods

### Membrane bioreactors

Three lab-scale Membrane Bioreactors (MBRs) were set up in this study. The reactors had a volume of 1 L and were filled with 400 mL medium (Fig. 1). A modified OECD degradation medium was used (https://www.oecd.org/chemicalsafety/testing/43735667.pdfhttps://www.oecd.org/chemicalsafety/testing/43735667.pdf). It contained 0.08 g/L peptone, 0.05 g/L meat extract, 0.015 g/L urea, 0.0035 g/L NaCl, 0.002 g/L CaCl_2_ x 2 H_2_O, 0.0001 g/L MgSO_4_ x 7 H_2_O and 0.0014 g/L K_2_HPO_4_. The pH of the medium was set to 7.5. The MBRs were constantly aerated using compressed air (pressure 0.5 bar, net O_2_ concentration 20 %) and a 2 cm stirrer was used for homogenization (600 rpm). A membrane-holder made of steel with two ultrafiltration membranes (pore size of 0.08 µm) of a total membrane area of 30 cm^2^ (Fig 1 a and b). The flow rate of influent and effluent was set at 10.5 mL/h, and MBRs were running continuously for 10 weeks. A backwash was performed weekly for 10 minutes to remove membranes’ cake layer to avoid membrane fouling. The hydraulic retention time of medium in the system was 38 h. The sludge retention time in the used MBR was infinite because no biomass was removed as excess sludge, except for sampling times.

**Fig. 1.**
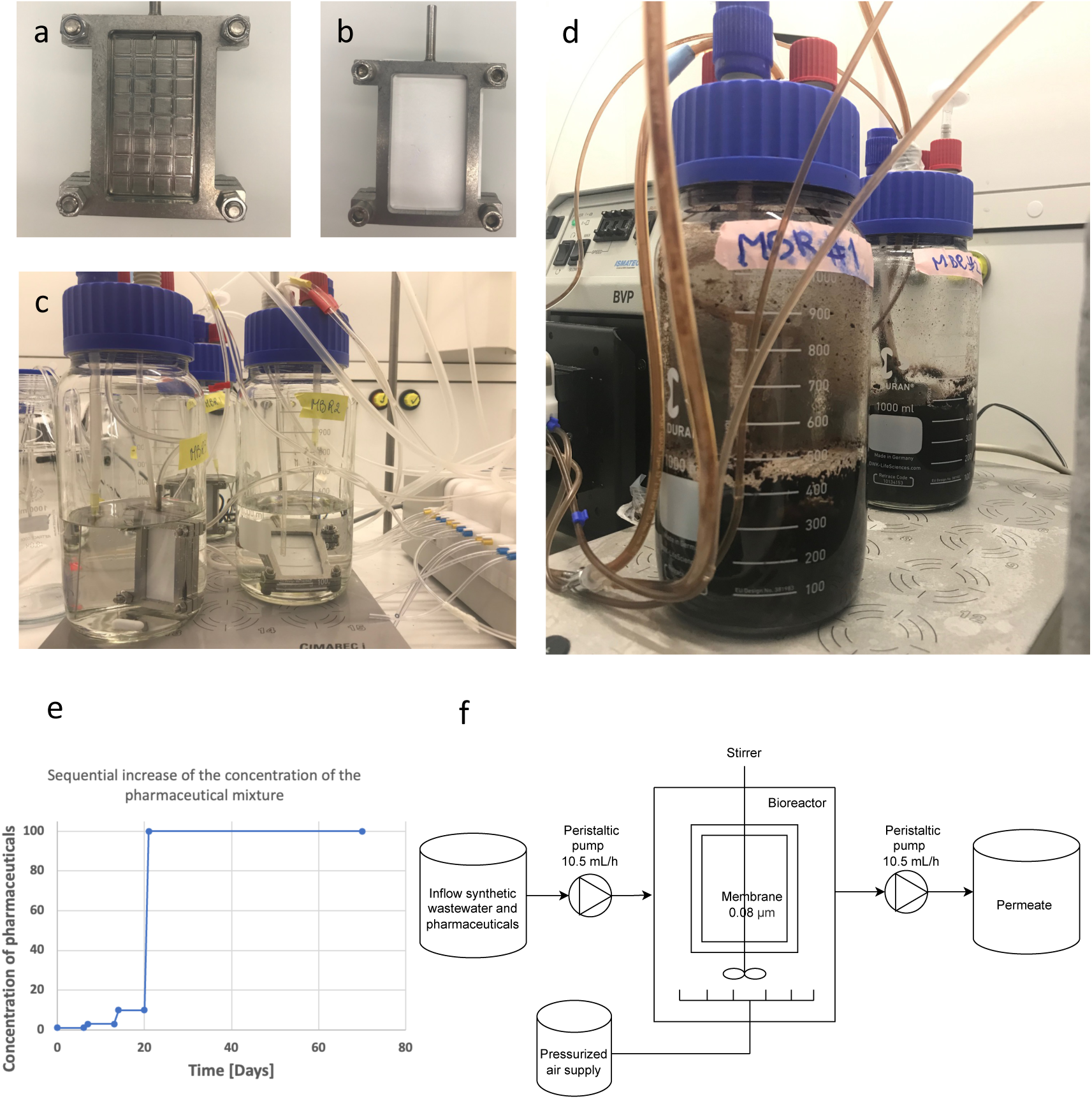
Technical setup of the MBRs. **(a)** Membrane-holder (steel) for placing two membranes (front side shown without membrane). The permeate hose was connected to the membrane-holder on the top. **(b)** Ultrafilter-membrane (0.08 µm) placed in in the membrane-holder. **(c)** Overview of MBRs set up before inoculation: membrane-holder, air spargers, magnetic stirrer for homogenization. **(d)** MBR 1 and 2 on day 21 of incubation with pharmaceuticals. **(e)** Pharmaceutical concentration gradient applied to the influent of MBR1 and MBR2. **(f)** Schematic overview of MBR settings. The hydraulic retention time of the system was 38 h.

Each MBR was inoculated with an activated sludge sample (1% v/v) of a WWTP. While one MBR was just operated with OECD degradation-medium as a control (MBR control), two MBRs were fed with the mixture of pharmaceuticals (MBR1 and MBR2). The pharmaceuticals used in this study were ibuprofen, diclofenac, enalapril, caffeine, atenolol and paracetamol and were all dissolved in the influent of MBR1 and MBR2. The starting concentration of pollutants was 1 mg/L for one week, afterwards the concentration of pollutants was weekly increased to 3 mg/L and 10 mg/L. Finally, after running of the MBRs for three weeks, the final concentration of pollutants was set at 100 mg/L and kept constant for another seven weeks.

### Sampling of the MBRs

5 mL of sample were taken from each of the three MBRs at different stages. Samples were taken directly from the bioreactor and not from the permeate. Samples were taken on the last day of incubating with 1 mg/L, 3 mg/L, 10 mg/L, respectively, and then taken weekly when pharmaceutical concentration was set at 100 mg/L. Furthermore, the original wastewater sample, which was used as inoculum, was analyzed by 16S rRNA sequencing. MBR_control (no pharmaceutical added) was sampled simultaneously to allow comparison with the MBRs exposed to the mixture of pharmaceuticals. The samples were centrifuged at 16,000 x g for 5 minutes. The pellet was used for DNA extraction, while the supernatant was used for HPLC analysis.

### Batch cultures with single micropollutant as substrate

Five batch cultures were set up (volume 100 mL of modified OECD-medium see above) with addition of 400 mg/L of a single pharmaceutical (ibuprofen, enalapril, caffeine, atenolol, paracetamol), respectively. One milliliter sample of MBR1, which was running for 9 weeks, was used as inoculum for each pharmaceutical. After seven days of growth, 1 mL of the batch culture was transferred again into fresh medium with the same conditions. Daily samples (1 mL) were taken for HPLC analysis to study the degradation potential of the microbial communities growing on each pharmaceutical, and 2 mL-samples were taken on day 3 for DNA extraction and sequencing.

### DNA extraction and sequencing

DNA was extracted using the ZymoBIOMICS DNA Miniprep Kit (ZymoResearch) by following the manufacturer’s instructions. The V4 region of the 16S rRNA gene were amplified and a DNA-library was made by using the Quick-16S™ Plus NGS Library Prep Kit (V4) (ZymoResearch). 4 pM DNA library (spiked in with 25 % PhiX) was sequenced in-house using Illumina MiSeq by following manufacturer’s instructions. Sequencing data were processed based on primer sequences, quality, error rates and chimeras using the r-package *dada2* (Callahan et al., 2016). The sequence table was aligned to the SILVA ribosomal RNA database (Quast et al., 2012), using version 138 (non-redundant dataset 99). A phyloseq object was created using the *phyloseq* r-package (McMurdie & Holmes, 2013), consisting of amplicon sequence variant (ASV) table, taxonomy table and sample data. For further analysis, the r-packages *phyloseq* (McMurdie & Holmes, 2013) and *vegan* (Oksanen et al., 2019) were used. The phyloseq object, metadata and the detailed R code for analysis are available on github (https://github.com/Marcel2907), and raw sequencing data are available on NCBI SRA SUB13057474.

### HPLC analysis

Pharmaceuticals were separated on a Hi-Plex Na column by high-performance liquid chromatography (HPLC) (Agilent Technologies) by applying a flow rate of 0.7 mL/min with water and methanol as mobile phase. The pharmaceuticals were detected using UV/VIS DAD detector. The mobile phase ratio started at 95:5 VV of, respectively, 0.1 % formic acid in Millipore water (A) and methanol (B). The B gradient was from 5% to 95% within 15 minutes and it allowed the analysis of all the six micropollutants in one run. The retention times were as follows: ibuprofen eluted at 12.16 minutes, diclofenac at 11.81 minutes, enalapril at 9.7 minutes, caffeine at 8.18 minute, atenolol at 6.21 minutes, paracetamol at 6.64 minutes. The detection wavelength was set at 230 nm for paracetamol, ibuprofen, atenolol, caffeine, diclofenac and at 205 nm for enalapril. A standard curve was generated for each pollutant (1 mg/L-1 g/L).

## Results

### Efficiency of microbial communities to remove multiple micropollutants within MBR1 and MBR2

Two MBRs (MBR1 and MBR2) fed constantly with synthetic wastewater contaminated by a mixture of six pharmaceuticals over a period of 10 weeks. After three weeks of incubation with 1 mg/L, 3 mg/L and 10 mg/L with all six pollutants, the concentration was changed to 100 mg/L and kept constant for 7 weeks. In week 7, 8, 9, and 10, the removal efficiency of each pollutant was analyzed by HPLC. Microbial communities within MBR1 and MBR2 evolved and became able to degrade most pollutants, except diclofenac, which concentration stayed stable in the MBRs (Fig. 2). However, the removal efficiency fluctuated between the different time points within MBR1 and MBR2. The microbial community within MBR1 was able to remove atenolol in a range of 55-100%, caffeine from 70-100%, enalapril from 34-100%, Ibuprofen in a range of 35-100%, and paracetamol from 71-100%. The evolved microbial community within MBR2 was able to remove atenolol in a range of 44-100%, caffeine from 67-100%, enalapril from 39-100%, Ibuprofen in a range of 0-100%, and paracetamol from 56- 100%. Ibuprofen removal is of interest because the performance of MBR2 to degrade it changed strongly from week 9 – week 10.

**Fig. 2.**
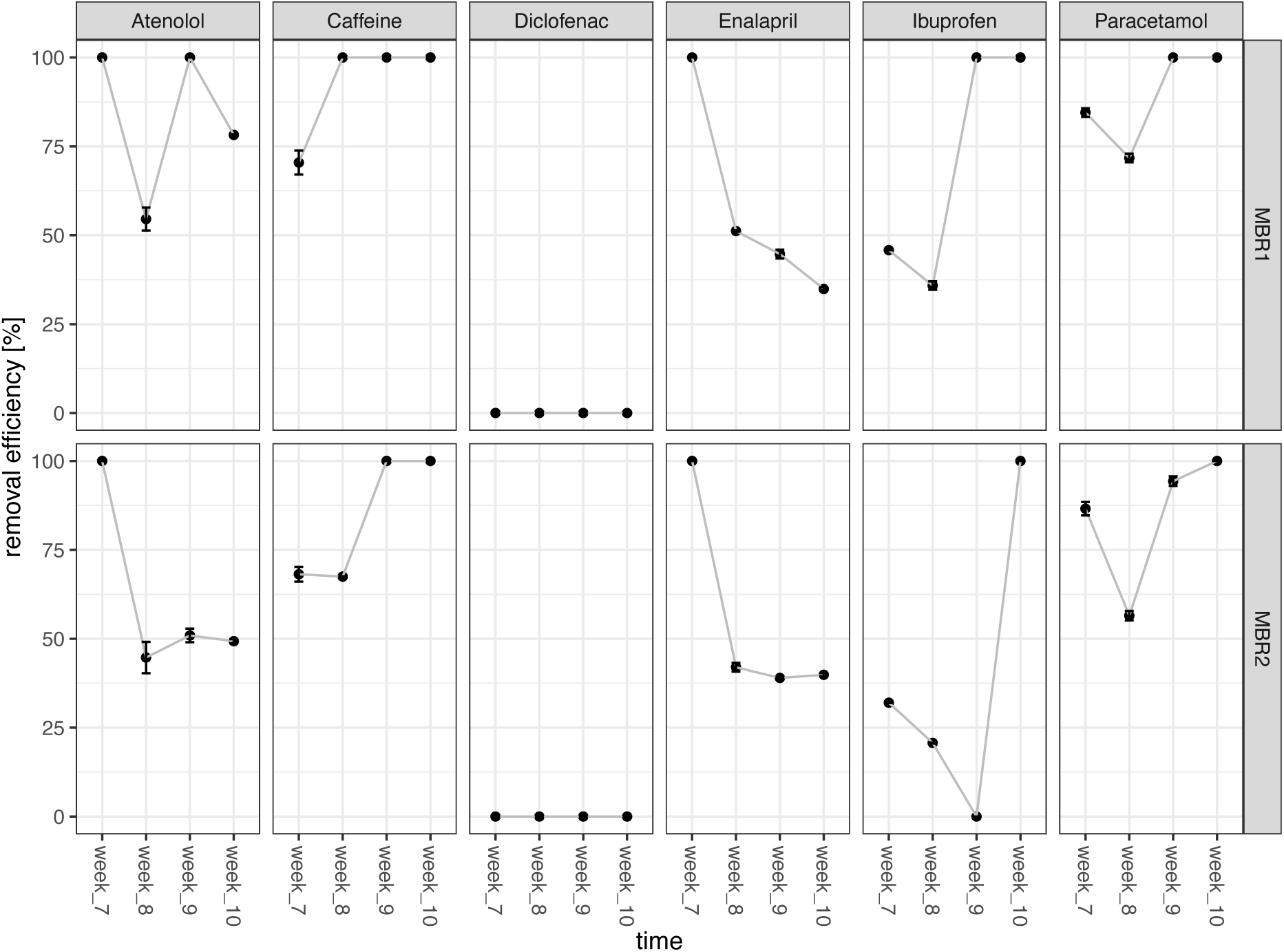
Efficiency of microbial communities to remove pharmaceuticals in MBR1 and MBR2 from week 7 to week 10 of cultivation. Influent-concentration of each pharmaceutical was 100 mg/L. Samples were taken ones per week directly from the bioreactor, and the concentration of pharmaceuticals in MBR1 and MBR2 was measured by means of HPLC and compared with influent concentration, to calculate the removal efficiency per reactor.

Interestingly, while the potential of degrading Ibuprofen, paracetamol and caffeine increased over time, the potential to degrade enalapril and atenolol decreased. No formation of degradation compounds of the micropollutants were detected via HPLC.

### Microbial community composition of MBR1, MBR2 and MBR_control

Microbial communities in pharmaceutical-treated MBR1 and MBR2 showed strong dynamic changes during application of the pharmaceutical gradient (1 mg/L-3mg/L-10mg/L-100mg/L) (Fig. 3 a and b). Once setting 100 mg/L of pollutant in the influent, the microbial communities of both MBRs became stable and reached a steady-state (Fig. 3, Fig. 4). Based on comparison with microbial community composition of MBR control, the microbial community of MBR1 and MBR2 was assumed to explain pharmaceutical degradation, since MBR1 and MBR2 communities differed significantly from the MBR_control. During incubations from week 7 – week 10 with 100 mg/L micropollutants, both MBR1 and MBR2 were dominated by stable microbial communities of *Achromobacter* (up to 39 %), *Cupriavidus* (up to 12 %), *Pseudomonas* (up to 17 %) and *Leucobacter* (up to 22 %), identifying these microbial genera as highly important for the degradation of highly concentrated pharmaceuticals (Fig 3a and b). By comparing the relative abundance of these microorganisms in MBR1 and MBR2, slight differences were detected based on sampling time and reactor (Fig. 3b). While *Achromobacter* and *Cupriavidus* increased their relative abundances with increasing pharmaceutical concentration, other bacterial members showed the opposite trend: *Comamonas* was highly abundant (up to 24%) at low concentration (0, 1, 3 and 10 mg/L) of pharmaceuticals, but vanished at 100 mg/L. Also, *Flectobacillus* showed interesting patterns, reaching very high relative abundances (up to 36 %) at 3 mg/L micropollutant concentration, while no longer present when the feed contained 100 mg/L of each pharmaceutical. *Acinetobacter*, *Leucobacter* and *Pseudomonas* were present during the whole experiment at various sample points and concentrations but differed in their relative abundances between MBR1 and MBR2: While the relative abundance of *Leucobacter* was higher in MBR1, *Acinetobacter* and *Pseudomonas* were more dominant in MBR2 (Fig. 3b).

**Fig. 3.**
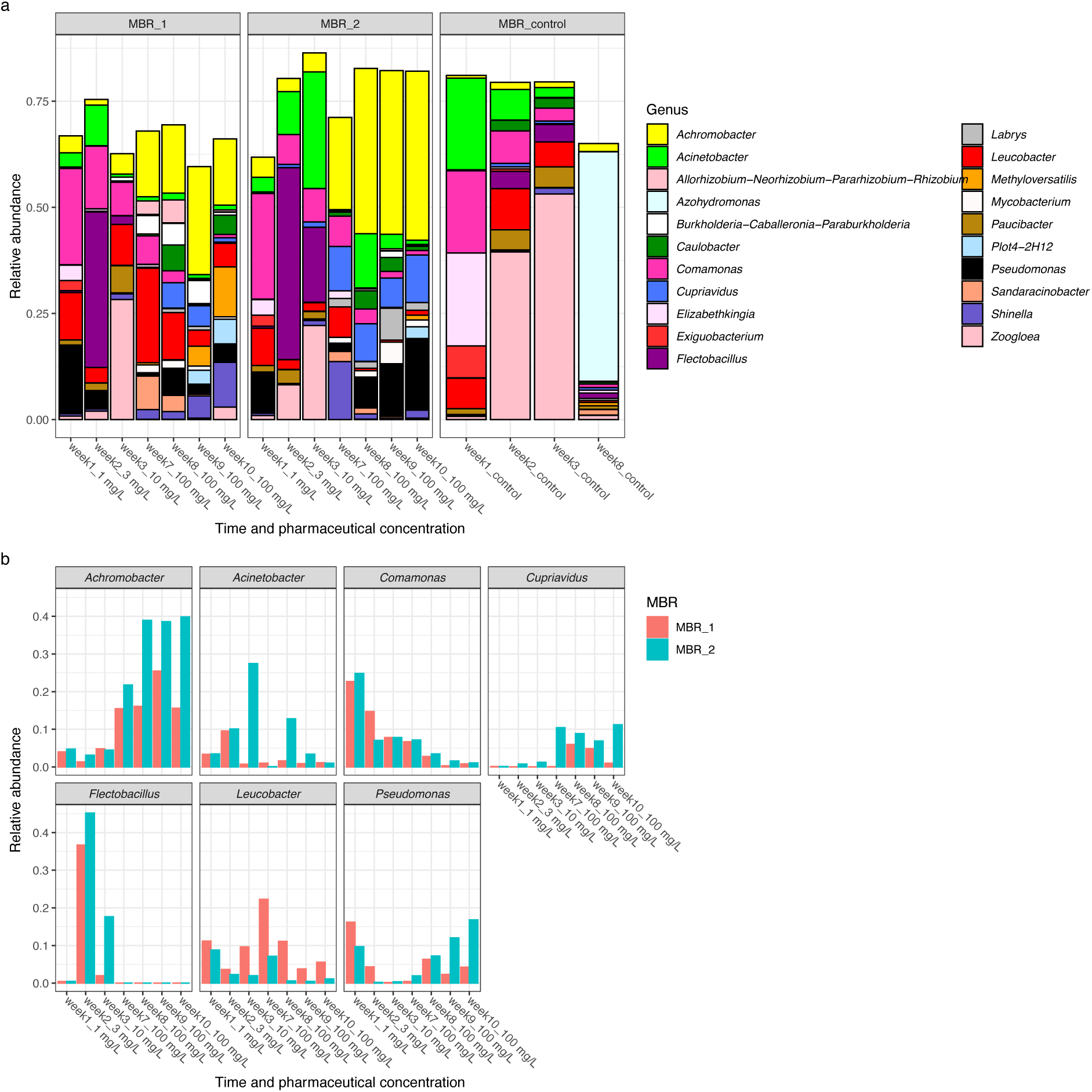
Microbial community composition of MBR1, MBR2 and control MBR. **(a)** Relative abundance on genus level for MBR1, MBR2 and control MBR. Genera with relative abundances > 5 % in at least one sample were included in the plot. **(b)** Relative abundances of most dominant genera of the pharmaceuticals treated MBR1 and MBR2 over time and pharmaceutical-concentration. The x axis shows the time and given concentration of pollutants.

**Fig. 4.**
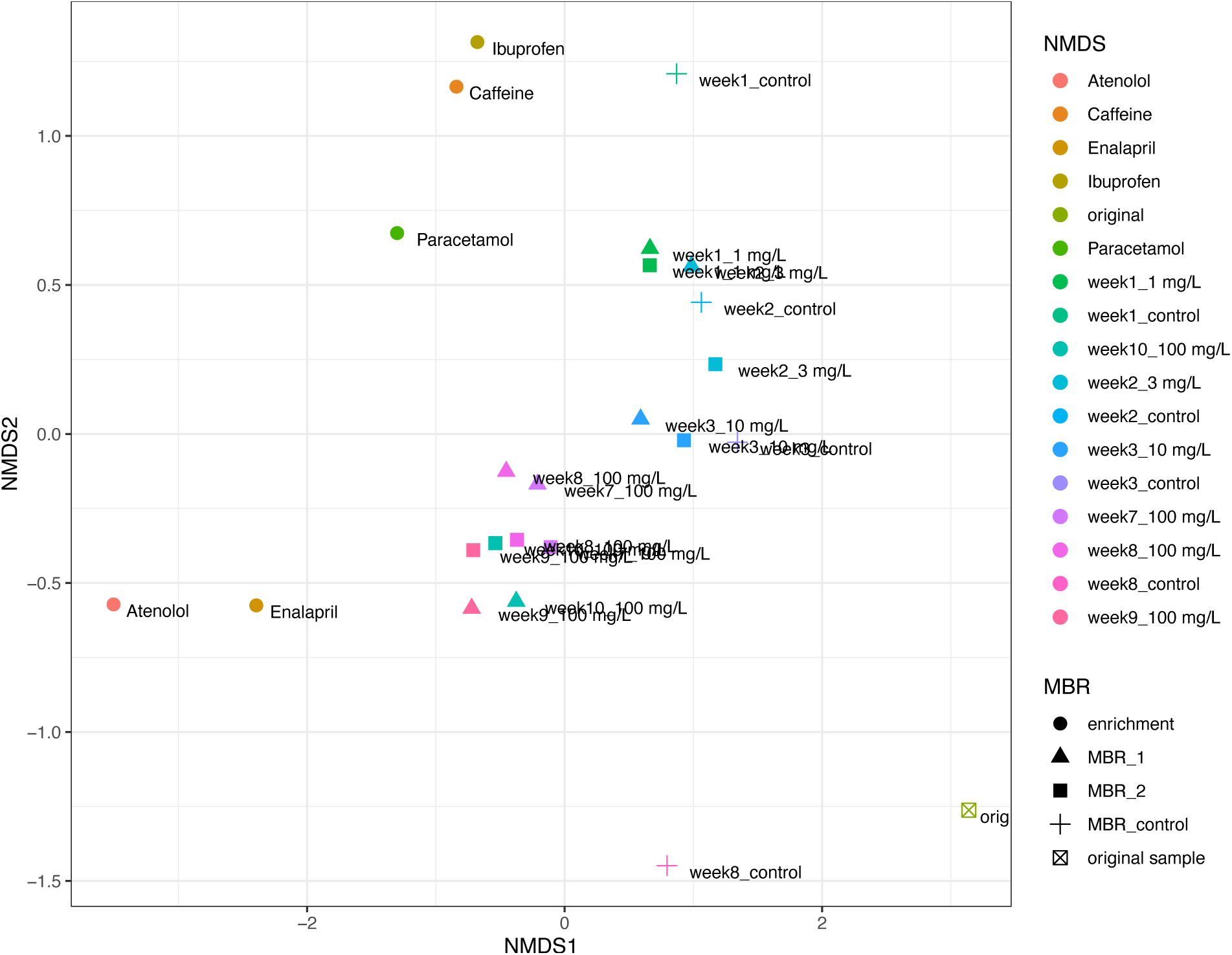
NMDS analysis based on Bray–Curtis distance of microbial communities from MBR1, MBR2, MBR control and the single micropollutant batch cultures. Distances of the microbial communities dependent on time points, pharmaceuticals (concentration and presence) are shown. The dots represent enrichment cultures grown on a single micropollutant as substrate. The triangles and the squares are the samples taken from MBR 1 and 2, respectively, at different time points and in presence of pharmaceutical. The crosses represent control MBR that was operated with OECD medium without pharmaceutical spike. The inoculum taken from WWTP is shown as a crossed square. Stress is 0.10.

NMDS analysis showed the dynamics of a developing microbial community under the increasing concentration of pharmaceuticals in the influent during first 3 weeks along the NMDS2 axis and demonstrated the stable state of the microbial communities from week 7 to week 10, clustering at the middle of the plot (Fig. 4). Furthermore, NMDS analysis demonstrated the strong distance of microbial communities within treated (MBR1 and MBR2) with untreated MBRs (MBR control). In addition, NMDS analysis highlighted that the adapted microbial communities of MBR1 and MBR2 differ strongly from the original activated sludge that was used as inoculum. While NMDS analysis (Fig.4) revealed a comparable microbial community in MBR1 and MBR2, slight differences in micropollutant removal efficiency of MBR1 and MBR2 were observed (Fig. 2), probably due to small differences on relative abundances of specific genera.

### Micropollutant degradation by microbial communities within the single-substrate batch cultures

The evolved microbial community of MBR1 (from week 9) was used as inoculum to prepare 5 batch cultures exposed each to single pollutant (with 400 mg/L pharmaceutical). The batch cultures were grown for four days, and the microbial community composition and pollutant concentration was determined (Fig. 5). Results indicated that all five substrates were (partly) removed, however, with very variable efficiencies. The microbial community of the culture incubated with paracetamol was able to degrade the entire 400 mg/L of the substrate, and the microbial community consisted mainly of *Sphingobacterium*, *Pseudomonas* and *Achromobacter*. The microbial community exposed to caffeine was also able to remove 400 mg/L caffeine, and microorganisms detected in high relative abundance were *Acinetobacter*, *Sphingobacterium*, *Pseudomonas* and *Chryseobacterium*. A comparable composition of microorganisms was found in the batch culture incubated with ibuprofen, and this consortium was able to remove 85 mg/L of the pollutant within 4 days. The batch culture exposed to atenolol was able to remove 57 mg/L atenolol, and the community consisted mainly of *Sphingomonas*, *Paucibacter* and *Burkholderia*. The batch culture exposed to enalapril was able to remove 35 mg/L enalapril and consisted mainly of *Klebsiella* and *Burkholderia*. The results of the microbial communities incubated with single substrates showed different microbial community compositions. The pharmaceutical substrate influenced the microbial community composition. Moreover, the microbial community of batch cultures exposed to a single pharmaceutical differed from the microbial communities found in MBRs as shown by NMDS (Fig. 4).

**Fig. 5.**
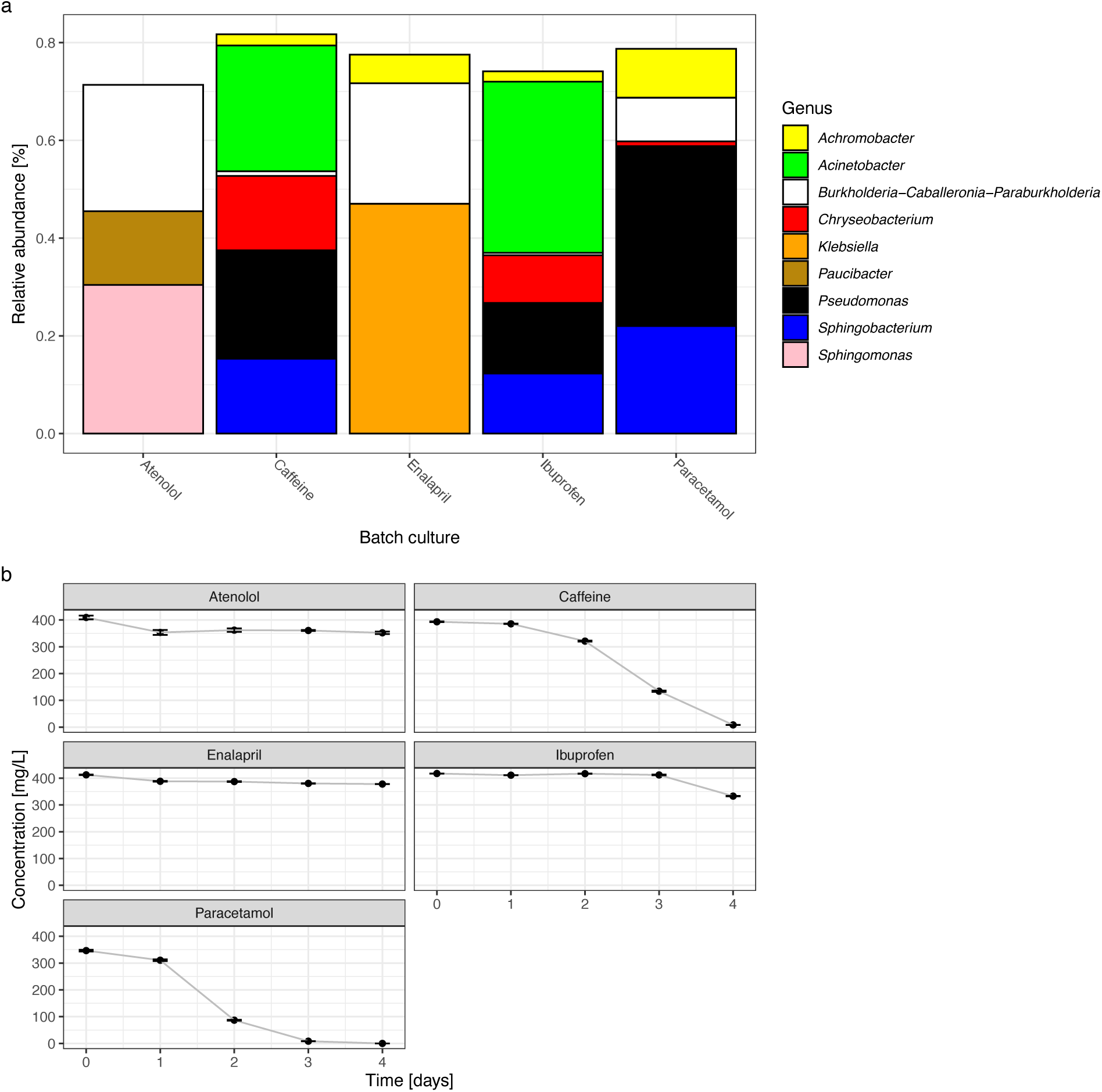
Microbial community composition and pharmaceutical degradation in batch cultures treated with single pharmaceuticals. **(a)** Relative abundance on genus level in the batch cultures grown for 4 days on single pharmaceutical. Sample from MBR1 were used (week 9) as inoculum. Genera with relative abundances > 10 % in at least one sample were included in the plot to identify key players of the respective culture. **(b)** Concentration of pharmaceuticals [mg/L] in the batch cultures (n=3).

## Discussion

The performed study gave first insights into the removal efficiency of a mixture containing six pharmaceuticals by microbial communities of lab-scale MBRs. The microbial communities evolved in the pharmaceuticals treated MBRs and showed constant and strong removal efficiencies of 50-100 % for caffeine, atenolol and paracetamol (50-100 %). Since these three pharmaceuticals are highly accumulating in WWTP with increasing concentrations in current and future scenarios, it is important to analyze their biodegradation potential. Caffeine is of particular interest, since this compound is one of the most concentrated pollutants found in WWTP, reaching already mg/L concentration (Li et al., 2020). As reported in several studies, our data confirm that the removal efficiency for caffeine by microbial communities is high (Li et al., 2020; Shanmugam et al., 2021; Summers et al., 2015). During the last years, biodegradation of atenolol, one of the most consumed beta-blockers worldwide, was in the focus of several studies and novel microbial degradation pathways of this compound were reported (Yi et al., 2022). Our data suggest that under the given hydraulic retention time of 38 h, the removal efficiency by the microbial communities was high, and comparable to previous studies focusing on the biodegradation of atenolol (Rezaei et al., 2022). The removal efficiency of 100 mg/L paracetamol was very high in both MBRs (100 % in MBR1, 95 % in MBR2). This confirms the microbial tendency of using paracetamol as substrate for microbial metabolism (Żur et al., 2018), especially by *Pseudomonas* strains (Rios-Miguel et al., 2022) that were also found in the operated MBRs of the present study.

Regarding ibuprofen, a pharmaceutical compound of high environmental concern (Chopra & Kumar, 2020), the removal efficiency of MBR1 increased over time to 100%. This demonstrates the potential of adaptation and evolutionary processes to improve the removal capacity for specific pollutants in MBR-operating systems (Hoinkis et al., 2012). Interestingly, MBR2 showed strong fluctuating removal efficiencies of ibuprofen per week, ranging from 0- 100%. Nevertheless, after 10 weeks of operation of MBR2, the microbial community was also able to remove 100 % of ibuprofen. Since no significant changes of the microbial community can be observed within these time points, varying abiotic parameters like oxygen intake and pH could explain these fluctuations and will be observed in future studies.

Biodegradation of ACE inhibitors, used in form of enalapril, is still poorly understood compared to the other micropollutants of this study. The removal efficiency under the given hydraulic retention time was around 30% at the end of the experiment, and interestingly showed a decreasing removal efficiency over time.

Diclofenac, a drug with high persistence in the environment (Sathishkumar et al., 2020), was the only micropollutant that was not removed by the evolved microbial communities at all, which demonstrates the need of focusing on the bioremediation of this compound. Interestingly, the microbial community found in the MBRs contained known diclofenac-degrading genera like *Labrys* (Moreira et al., 2018) in a relative abundance up to 7 %. However, no removal of diclofenac was observed. This could be explained by the fact that the microbial communities within the MBRs are confronted to multiple substrates with easier biodegradability than diclofenac. This demonstrates the high complexity of dealing with multiple substrates (like in real scenarios) instead of just one driver (Suleiman et al., 2022). The given concentrations of 100 mg/L of each pollutant are significantly higher compared to natural concentrations found in wastewater, but such high concentrations were necessary to allow microbes to grow on these compounds for achieving biomass. Furthermore, as the concentration of micropollutants in wastewater is constantly increasing, it is essential to assess their impact on microbial communities.

Our results are identifying potential key players of microbial communities, namely *Achromobacter*, *Cupriavidus*, *Pseudomonas* and *Leucobacter,* for simultaneous removal of multiple combined micropollutants. Recent studies showed that *Achromobacter* was associated with the bioremediation of pharmaceuticals, especially the antibiotic sulfamethoxazole (Liang & Hu, 2021). Various *Pseudomonas* species were associated with ibuprofen and paracetamol degradation (Rios-Miguel et al., 2022; Rutere et al., 2020), and the genus *Cupriavidus* was reported to degrade polluting aromatic compounds (Pérez-Pantoja et al., 2008). Therefore, while these genera were already associated with degradation of pharmaceutical micropollutants, it is to our knowledge the first time that these genera were found in one stable and active microbial community dealing with multiple pharmaceutical pollution.

Our data demonstrated changing microbial communities during the gradually increasing pharmaceuticals concentrations in the synthetic influent, which demonstrates that pharmaceuticals concentrations affect the dynamics and compositions of microorganisms. One major question is if the established microbial community, which is adapted to high pharmaceutical concentrations, is still able to degrade *in situ* pollutants in a µg/L scale. The results of this study are indicating a change of key players in batch cultures on single micropollutants, which demonstrates that multiple pollutants exposure affects microbial communities differently compared to cultures exposed to a single pharmaceutical. Besides the key players that were already identified in the MBRs, the batch cultures were, dependent on the substrate, dominated by *Acinetobacter*, *Sphingomonas* or *Sphingobacterium*. *Acinetobacter* were dominant in cultures incubated with caffeine and ibuprofen, respectively, and members of this genus were already reported to show good efficiency in degrading crude oil (Zhang et al., 2021). *Sphingomonas* was dominant in culture with atenolol, which was partly degraded, and was already associated with degradation of ibuprofen in recent studies (Murdoch & Hay, 2013). *Sphingobacterium* was dominant in cultures with caffeine, paracetamol, and ibuprofen, and was already reported to degrade complex compounds such as 17α-ethynylestradiol (Haiyan et al., 2007). Interestingly, these three genera had a high relative abundance in the batch cultures with single micropollutant, but not in the MBRs fed with the mixture of all pharmaceuticals. The critical trait of wastewater is the complex mixture of a multitude of compounds present at trace and high concentrations. Therefore, more studies focusing on multiple micropollutant are needed, since our study suggests that microbial communities and their degradation potential of pharmaceuticals varies, depending on single or multiple exposure to pharmaceuticals and their concentration.

## Conclusion

The issue of water remediation is needed due to the predicted increase in pharmaceutical consumption and the increasing demand for higher removal of pollutants in treated water. The six pharmaceuticals used in this study are found in high concentrations in influents and effluents of wastewater plant, with an increasing trend. The adaptive laboratory evolution in this study showed that after a prolonged time under pharmaceuticals concentration gradient pressure, the microbial community reached a stable state at 100 mg/L pharmaceuticals exposure. The communities of the two MBRs were able to degrade ibuprofen, paracetamol, caffeine, enalapril and caffeine with a fluctuating but strong efficiency. These fluctuations will be analyzed in future studies by controlling oxygen intake, pH and temperature of the operating system. The communities evolved in MBR1 and MBR2 after 10 weeks of incubation differed significantly from MBR control (and inoculation sample), proving that the microbial communities adapted successfully to the pharmaceuticals as substrate for their subsistence.

Furthermore, it was possible to identify specific species as potential key players for the degradation of single highly concentrated pharmaceuticals. Promising candidates for removing pharmaceutical micropollutants in WWTP such as *Achromobacter*, *Pseudomonas* and *Cupriavidus* were dominating in the late stage of the adaptation. This preliminary study can be further developed for real case application to wastewater treatment plant as polishing step for the removal of pharmaceuticals that are not efficiently removed in the biological step.

## Acknowledgements

This work was funded through the European Union’s Horizon 2020 project NMYPHE under grant ID 10106625.

The authors would like to thank Sebastian Hedwig, Jan Svojitka, Roman Schäfer and Patrik Eckert of the group of Prof. Michael Thomann (University of Applied Sciences and Arts Northwestern Switzerland, Institute of Ecopreneurship) for the support of constructing lab-scale MBRs.

